# Effect of soybean molasses-adsorbents on in vitro ruminal fermentation characteristics, milk production performance in lactating dairy cows

**DOI:** 10.1101/496224

**Authors:** Liang Chen, Bin Li, Ao Ren, Zhiwei Kong, Chuanshe Zhou, Zhiliang Tan, Shaoxun Tang, Rejun Fang

## Abstract

This study aimed to evaluate the *in vitro* fermentation characteristics of corncob powder (CRP), wheat bran (WB), rice husk (RH), defatted bran (DB) and soybean hulls (SH) when mixed with soybean molasses at a ratio of 30:100 (dry matter basis), using a batch culture incubation. During *in vitro* study, SH showed better fermentation characteristics (including greater maximum gas production, shorter time to achieve half of *Vf*, greater concentrations of acetate, propionate and total VFA, and less initial fractional rate of degradation) than other four substrates, while WB had the greatest values of DM and NDF disappearance, NH3-N and butyrate concentrations among substrates. A randomized complete block designed *in vivo* experiment was conducted with 24 Holstein cows (534 ± 58 kg BW, 2.8 ± 0.7 parity, 129 ± 23 d in milk) randomly assigned to three experimental diets: Control, WB (WB adsorbed to soybean molasses replaced 150 g of corn meal per 1000 g of diet dry matter [DM]) or SH (SH adsorbed to soybean molasses replaced 100 g of wheat bran and 50 g corn meal per 1000 g of diet DM). The results indicated that cows received WB diet had greater (*P*<0.01) milk fat and total milk solid content than cows fed control and SH diets, and cows received WB and SH diets tended to have greater (*P*<0.01) milk protein content and blood glutamic-pyruvic transaminase concentration than cows fed control diet. Furtherly, cows received WB diet had greater (*P*<0.01) blood amylase and lactate dehydrogenase concentration than that of cows fed control diet during middle lactation.

In conclusion, dietary supplementation of molasses adsorbed by-products like WB and SH have positive effect on promoting rumen fermentation, milk quality and blood metabolism in early- and middle-lactating dairy cows. The results offered a new products and feeding way in dairy farming

## Introduction

Soybean molasses is a by-product of soybean meal concentrate. The molasses byproduct results from the separation of solids and the evaporation of ethanol from the liquid fraction during the ethanol extraction processes of concentrated soybean meal. It is rich in oligosaccharides, saponins, isoflavones and other phytochemicals (Shi et al. 2013). Most of the carbohydrates can be fermented rapidly in the rumen by the microbes as energy sources, leading to efficient utilization of the rapidly degradable nitrogen fraction and greater microbial protein synthesis. The net result can be increased milk protein production.

Previous studies mainly focused on molasses extracted from sugarcane and beet, which have been widely used in the animal feed industry. Molasses is a sugar-containing liquid feed that can increase the ruminal fermentability of the diet, while stimulating dry matter intake (DMI) (Firkins et al., 2008). It also serves as a fat carrier in a diet and is used to enhance mixing of ingredients to prevent sorting (Murphy et al., 1997). Feeding a sugar-based product can change the ruminal fermentation pattern, decrease ruminal ammonia (NH_3_) concentration in dairy cows (Broderick and Radloff, 2004; Broderick et al., 2008), and increase ruminal butyrate concentration (Hristov and Ropp, 2003; DeFrain et al., 2006). It is well-known that sugars can be rapidly fermented in the rumen, theoretically leading to lactic acid production and a decline in ruminal pH, which potentially depresses fiber digestibility (Oelker et al., 2009). Martel et al. (2011) reported that dietary supplementation with cane molasses affected volatile fatty acid (VFA) concentration, milk production, and milk fat and protein yields. Furthermore, supplementation of blended molasses (50% beet sugar molasses and 50% yeast molasses) can alleviate the decrease of feed intake, and increase milk production and milk protein content in dairy cows during heat stress (Zhang, et al., 2013). Broderick and Radloff (2004) reported that replacing high-moisture corn with molasses improved fiber digestibility, likely reflecting a stimulatory effect of molasses on fiber-digesting ruminal bacteria.

Our first hypothesis for this study using a batch culture *in vitro* fermentation technique was that WB and SH had better *in vitro* fermentation characteristics among five different feeds adsorbed to soybean molasses. and our second hypothesis for this study using lactating dairy as the experimental animals was that WB and SH have positive effect on improving milk quality and promoting blood metabolism in lactating dairy cows.

## Material & Methods

The experiments were conducted according to the animal care guidelines of the Animal Care Committee, Institute of Subtropical Agriculture, The Chinese Academy of Sciences, Changsha City, Hunan Province, China (No. KYNEAAM-2006-0015).

### 1.1 In vitro experiment

#### 1.1.1 Fermentation substrates

Corncob powder (CRP), wheat bran (WB), rice husk (RH), defatted bran (DB) and soybean hulls (SH) were mixed with soybean molasses at a ratio of 100:30 (DM basis), dried at 65°C for 24 h, ground through a 1-mm sieve and stored in an airtight bag until further assays(Offered by Fengyi (Shanghai) biotechnology research and development center co. LTD. Shanghai, 200137, China). The chemical compositions of the five soybean molasses-adsorbents are listed in Table 1.

**Table 1.**
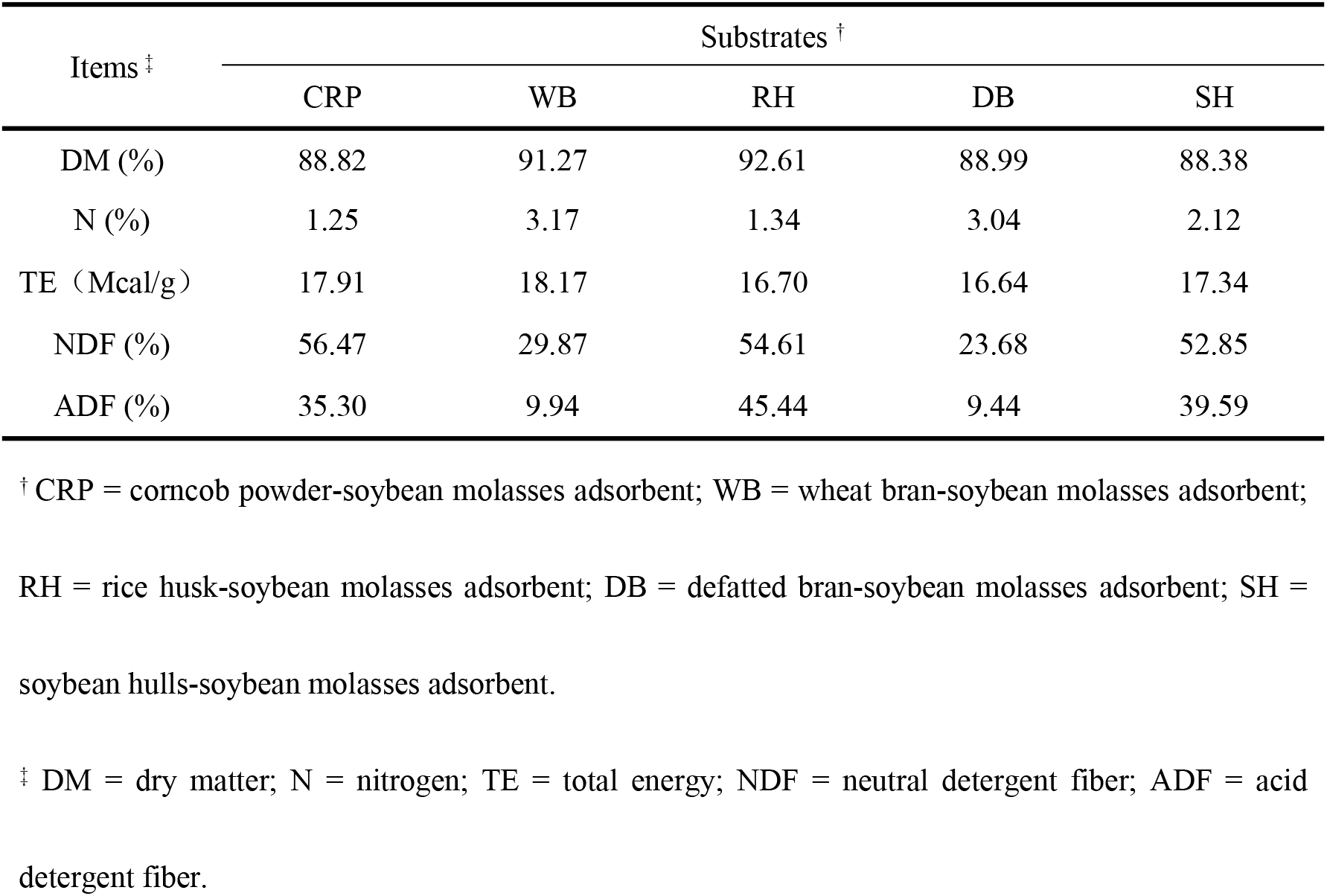
Chemical composition of five soybean molasses adsorbed substrates

#### 1.1.2 In vitro gas production and sampling

The *in vitro* study was designed as a single factor randomized block design to evaluate the effects of five molasses-adsorbents. *In vitro* batch culture solutions were prepared using macroelement solution, buffer and reducing agent. The buffer was prepared as described by Tang et al. (2006) and it was kept anaerobic by continuously pumping carbon dioxide for 2 h. Rumen fluid was obtained from three rumen-cannulated Holstein dairy cows fed *ad libitum* a mixed diet of rice straw and concentrate (60:40, wt/wt). The diets were offered twice daily at 0500 and 1600 h. Rumen contents of each dairy cow were obtained from various locations within the rumen immediately before the morning feeding, mixed and strained through four layers of cheesecloth under a continuous CO_2_ stream. The obtained rumen fluid was then anaerobically combined with *In vitro* batch culture solutions in the proportion of 1 to 9 at 39°C.

A 1000 ± 3 mg sample of each substrate was accurately weighed into a 100 mL fermentation bottle (Wanhong Glass Instrument Factory, China) pre-warmed at 39°C, then 50 ml of the mixed fluids (rumen fluids:artificial saliva = 1:9, V/V) were dispensed into each bottle. Each sample was replicated three times at each incubation time point. Bottles containing only mixed fluids were incubated as blanks together with the bottles containing different molasses-adsorbents. All fermentation bottles were connected with pressure sensors (CYG130-12, SQ sensor, China) and incubated at 39°C. The pressure in all the bottles was recorded at 0, 1, 2, 4, 6, 12, 24, and 48 h during the process of *in vitro* fermentation. Three bottles for each treatment were removed from the incubator to stop the incubation and the pH of the fluid in each bottle was determined immediately. The undegraded residues were filtered through 2 layers of nylon cloth (40-µm pore size). The incubation fluid was sampled at 12, 24 and 48 h for determination of NH_3_-N and VFA concentrations.

### 1.2 In vivo experiment

#### 1.2.1 Experimental diets and design

A randomized complete block design was with 24 multiparous Holstein cows (534 ± 58 kg BW, 2.8 ± 0.7 parity) blocked into 8 blocks to ensure equal numbers of early-lactation (0-100 d) and mid-lactation (100-200 d) cows for each treatment. One cow per group was randomly assigned to one of three treatments：Control (basal diet); WB (WB adsorbed to soybean molasses replaced 150 g of corn meal per 1000 g of diet dry matter [DM]) or SH (SH adsorbed to soybean molasses replaced 100 g of wheat bran and 50 g corn meal per 1000 g of diet DM). The three experimental diets were formulated to meet the nutrient requirements of lactating cows according to NRC (2001), and the treatmetns were chosen based on the *in vitro* fermentation results (Table 2).

**Table 2.** Ingredients and chemical composition of experimental diets

| Ingredient | Group |  |  |
| --- | --- | --- | --- |
|  | CG | SH <sup>†</sup> | WB <sup>‡</sup> |
|  | Feeding (kg/d • cow) |  |  |
| Rice straw | 6.0 | 6.0 | 6.0 |
| Beet pulp | 3.0 | 3.0 | 3.0 |
| DGS<sup>§</sup> | 3.0 | 3.0 | 3.0 |
| Concentrate | 4.0 | 4.0 | 4.0 |
|  | (% of DM) |  |  |
| Corn meal | 43.1 | 33.1 | 43.1 |
| Soybean meal | 10.0 | 10.0 | 10.0 |
| Wheat bran | 18.0 | 13.0 | 3.0 |
| DGS<sup>§</sup> | 21.0 | 21.0 | 21.0 |
| SH adsorbed molasses | - | 15.0 | - |
| WB adsorbed molasses | - | - | 15.0 |
| CaHPO<sub>4</sub> | 1.5 | 1.5 | 1.5 |
| CaCO<sub>3</sub> | 1.3 | 1.3 | 1.3 |
| NaHCO<sub>3</sub> | 0.6 | 0.6 | 0.6 |
| NaCl | 0.5 | 0.5 | 0.5 |
| Premix<sup>¶</sup> | 4.0 | 4.0 | 4.0 |
| Chemical composition of concentrate (% of concentrate DM) |  |  |  |
| Dry matter | 93.98 | 94.11 | 94.63 |
| Crude protein | 18.89 | 19.66 | 19.13 |
| Calcium | 1.90 | 1.86 | 1.89 |
| Phosphorus | 0.80 | 0.77 | 0.79 |
| Neutral detergent fiber | 18.70 | 17.22 | 18.03 |
| RDP, % of CP | 59.81 | 60.14 | 59.29 |

<sup>†</sup> SH = soybean hulls-soybean molasses adsorbents.
<sup>‡</sup>WB = wheat bran-soybean molasses adsorbents.
<sup>§</sup>DGS = distillers grains with solubles.
<sup>¶</sup> Premix ( /kg) : 113.85g MgSO<sub>4</sub>•H<sub>2</sub>O, 2.69 g FeSO<sub>4</sub>•7H<sub>2</sub>O, 2.55 g CuSO<sub>4</sub>•5H<sub>2</sub>O, 9.54 g
$\text{MnSO}_4 \cdot \text{H}_2\text{O}$ , 9.60 g $\text{ZnSO}_4 \cdot \text{H}_2\text{O}$ , 30 mg $\text{Na}_2\text{SeO}_3$ , 60 mg KI, 180 mg $\text{CoCl}_2 \cdot 6\text{H}_2\text{O}$ , 500,000 IU
Vitamin A, 60 kIU Vitamin D, 2000 IU Vitamin E.

The experiment lasted 5 weeks. Throughout the trial, cows were housed in a tie-stall facility. Diets were offered adlibitum twice daily at 0500 and 1600 h, and had free access to clean water. Before starting the experiment, all cows were fed the same diets for 2-wk.

### 1.3 Sample collection and handling

The experimental diets were offered twice daily, the orts were collected and recorded once daily. Weekly composites of the concentrates, rice straw, orts, **DGS** (distillers grains with solubles) and beet pulp were obtained from daily samples of about 0.5 kg and stored at −20°C until analysis. Cows were milked twice daily, and individual milk yield was recorded at each milking during 5-weeks-expreriment. Milk samples were collected at 2 consecutive (p.m. and a.m.) milkings midway through wk 5 of the experimental phase for conventional analysis. Concentrations and yields of fat, protein, lactose, total solids (**TS**) and solids-not-fat (**SNF**) were computed as the weighted means from p.m. and a.m. milk yields on each test day. Blood samples were collected on the last day of wk 5 at 0500, 0700 and 1100 h, respectively. Ten mL of blood samples were collected every point-in-time from the coccygeal vein into Vacutainer tubes which included anticoagulation (heparin sodium). After sampling, tubes were kept on ice and immediately transported to the laboratory for centrifugation at 4000 × g for 10 min at 4°C, and plasma was stored at −80°C until assayed. 1.4 Chemical analyses

### 1.4 Chemical analyses

The DM and CP of *in vitro* fermentation substrates, concentrates, forage, orts, DGS and beet pulp were analyzed using the procedures of the Association of Official Analytical Chemists (AOAC, 2002). The NDF and ADF contents of the samples were determined using a Fibretherm Fiber Analyzer (Gerhardt, Bonn, Germany) according to Van Soest et al. (1991) with addition of sodium sulphite and alpha-amylase in the NDF analysis. The filtered residues were dried at 105°C for 12 h and weighed for *in vitro* DM disappearance (**IVDMD**) determination. The NDF contents of the dried residues were determined to calculate *in vitro* NDF (**IVNDFD)**. Total groos energy (**TE**) content was determined by an isothermal automatic calorimeter (5E-AC8018, Changsha Kaiyuan Instruments Co., Ltd, China) The NH_3_-N and VFA concentration was determined according to Chen et al (2017), Milk samples were analyzed for fat, protein, lactose, SNF and TS by infrared methods (Foss North America, Eden Prairie, MN; Ag-Source, Verona, WI).Glutamic-pyruvic transaminase (**GPT**), plasma ammonia (**AMM**), amylase (**AMY**), cholesterol (**CHO**), glucose (**GLU**), lactate dehydrogenase (**LDH**), triglyceride (**TG**), total protein (**TP**), and urea nitrogen (**UN**) were analyzed by kits (Beijing Leadman Biochemical Co., Ltd, Beijing, China) using auto-biochemical analyzer (Beckman CX4, Beckman Coulter, Inc. USA).

#### Calculation and Statistical Analysis

During the initial stages of the *in vitro* experiment, the correlation between the pressure in fermentation bottles and gas volumes was measured at 39°C, and the regression equation was then established:

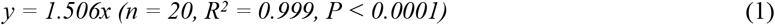

Where *y* represents gas volume (ml), *x* is the pressure in bottle (kPa), and 1.506 is a constant. The measured pressure was then converted to gas production (ml). *In vitro* gas production (GP) at 0, 1, 2, 4, 6, 12, 24, and 48 h were fitted to a logistic-exponential equation (Wang et al. 2011):

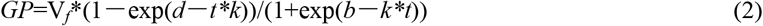

Where *GP* represents gas production at *t* time, *Vf* is the maximum gas production (ml), *k* represents gas production fraction (/h), *b* and *d* represent the shape of the gas production curve. The time (*t_0.5_*, h) when half of the maximum gas production was achieved and the initial fractional rate of degradation (*FRD_0_*, /h) were respectively calculated by employing the following two equations (Wang et al. 2011; Wang et al. 2013):

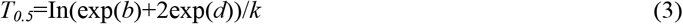

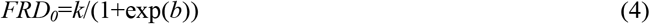

The GP, IVDMD and IVNDFD were corrected by subtracting the values obtained for the blanks. Data were analyzed by two-way ANOVA using the MIXED procedure of SAS (2001), and the incubation time was treat as a repeated factor. Results of milk production, milk quality and blood parameters were statistically analyzed using ANOVA and the MIXED procedure of SAS (2001). Duncan’s multiple range tests were used to compare differences among the three treatments. A P-value < 0.05 indicated statistical significance.

## 2 Results

### 2.1 In vitro experiment

#### 2.1.1 In vitro gas production characteristics of different molasses-adsorbents

The maximum gas production (*V_f_*) and *t_0.5_* of SH were both greater (*P* < 0.01) than that of CRP, WB, RH and DB, while no significant differences (*P* > 0.05) were observed among the other four molasses-adsorbents (Table 3). However, the *FRD_0_* (0.022 mL·h^−1^) of SH was the least among all molasses-adsorbents, and it was less than that of WB, RH and DB.

**Table 3.**
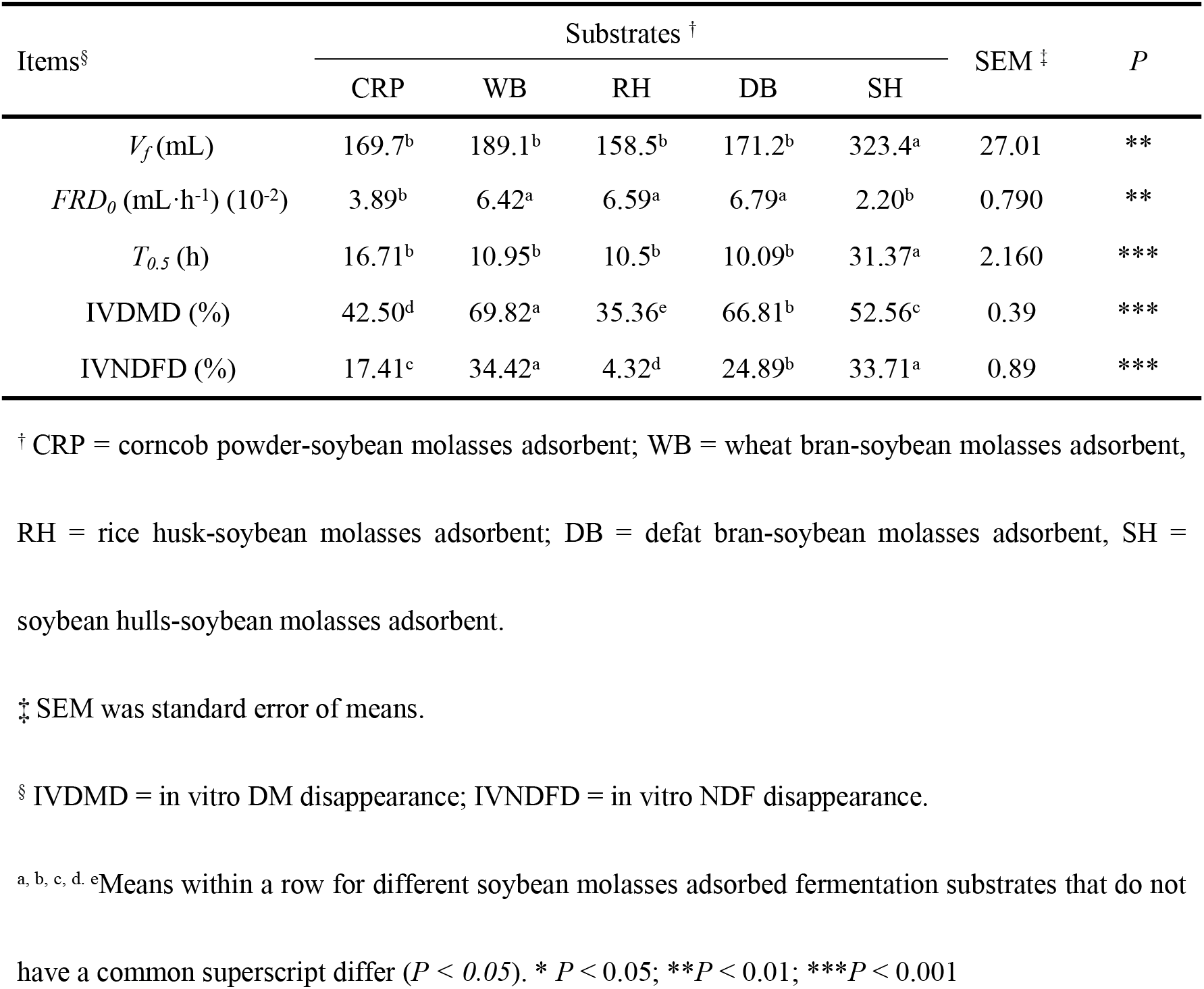
Effects of different soybean molasses adsorbed substrates on *in vitro* gas production parameters, IVDMD and IVNDFD

#### 2.1.2 IVDMD, and IVNDFD of different molasses-adsorbents

Differences (*P* < 0.0001) in IVDMD among the five molasses-adsorbents were observed (Table 3), with the IVDMD observed for WB (69.82%) being 27.3, 34.5, 3.0 and 17.3% greater than that of CRP, RH, DB, and SH, respectively. The IVNDFD of WB and SH were greater (*P* < 0.0001) than that of other three molasses-adsorbents, with the lowest IVNDFD observed for RH (4.32%).

#### 2.1.3 pH and NH_3_-N concentration of *in vitro* incubation fluids for different molasses-adsorbents

The range of pH values of the *in* v*itro* fermentation fluids was 5.89 to 6.75. The lowest pH value was for WB, with it being less (*P* < 0.0001) than that of the other four molasses-adsorbents (Table 4). The greatest NH_3_-N concentration (35.2 mg/dL) was obtained for WB, with it being greater (*P* < 0.0001) than that of the other four molasses-adsorbents.

**Table 4.** Effects of different soybean molasses adsorbed substrates on *in vitro* fermentation pH, NH_3_-N concentration, and VFAs concentration

| Items | Substrates <sup>†</sup> |  |  |  |  | SEM <sup>‡</sup> | P |
| --- | --- | --- | --- | --- | --- | --- | --- |
|  | CRP | WB | RH | DB | SH |  |  |
| pH | 6.34<sup>b</sup> | 5.89<sup>d</sup> | 6.75<sup>a</sup> | 6.01<sup>c</sup> | 6.00<sup>c</sup> | 0.03 | *** |
| NH<sub>3</sub>-N (mg/dL) | 5.09<sup>d</sup> | 35.18<sup>a</sup> | 15.91<sup>b</sup> | 12.95<sup>c</sup> | 12.68<sup>c</sup> | 0.35 | *** |
| Acetate (mmol/L) | 32.07<sup>c</sup> | 37.10<sup>b</sup> | 32.98<sup>bc</sup> | 35.73<sup>bc</sup> | 45.29<sup>a</sup> | 1.59 | *** |
| Propionate (mmol/L) | 19.44<sup>b</sup> | 25.41<sup>a</sup> | 19.83<sup>b</sup> | 24.81<sup>a</sup> | 27.08<sup>a</sup> | 1.45 | ** |
| Butyrate (mmol/L) | 4.14<sup>c</sup> | 9.42<sup>a</sup> | 6.72<sup>b</sup> | 7.65<sup>b</sup> | 7.47<sup>b</sup> | 0.53 | *** |
| TVFA (mmol/L) <sup>§</sup> | 56.65<sup>c</sup> | 75.99<sup>ab</sup> | 61.24<sup>c</sup> | 69.88<sup>b</sup> | 81.99<sup>a</sup> | 2.98 | *** |

<sup>†</sup> CRP = corncob powder-soybean molasses adsorbent; WB = wheat bran-soybean molasses adsorbent; RH = rice husk-soybean molasses adsorbent; DB = defat bran-soybean molasses adsorbent; SH = soybean hulls-soybean molasses adsorbent.
<sup>‡</sup> SEM was standard error of means.
<sup>§</sup> TVFA = total VFA.
<sup>a, b, c, d</sup> Means within a row for different soybean molasses adsorbed fermentation substrates that do not have a common superscript differ ( $P < 0.05$ ). \* $P < 0.05$ ; \*\* $P < 0.01$ ; \*\*\* $P < 0.001$

#### 2.1.4 VFA content of in vitro incubation fluids for different molasses-adsorbents

Acetate content of SH was greater (*P* < 0.0001) than that of the other four molasses-adsorbents (Table 4). The propionate content of SH was the greatest among the five molasses-adsorbents, with it being greater than (*P* < 0.01) that of CRP and RH. Butyrate content of WB was 56.1, 28.7, 18.8 and 20.7% greater (*P* < 0.0001) than that of CRP, RH, DB and SH, respectively. The SH and WB also had the greatest content of TVFA (*P < 0.0001*). There were no differences in A:P (*P > 0.05*) for all five molasses-adsorbents.

### 2.2 In vivo experiment

#### 2.2.1 Milk performance

The milk yield was 25.0 and 17.0 kg during early- and mid-lactation, respectively, and there were no differences (*P* > 0.05) among the three treatments for either lactation period (Table 5). The contents of lactose and SNF were not affected (*P* > 0.05) by the replacement of molasses-adsorbents. The milk fat and total solids contents of cows fed the WB treatment were greater (*P* < 0.01) than those fed control and SH treatments in early lactation, while there were no treatment differences (*P* > 0.05) in mid-lactation. The milk protein content of cows fed the CG treatment decreased by 0.34%, 0.20% and 0.17%, 0.16% (*P* < 0.01) compared with that of WB and SH treatments in early- and mid-lactation periods, respectively.

**Table 5.** Effect of different soybean molasses adsorbed substrates on milking performance in different lactating period in dairy cows

| Items | Group <sup>†</sup> |  |  | SEM <sup>‡</sup> | *P* <sup>§</sup> |
| --- | --- | --- | --- | --- | --- |
|  | CG | WB | SH |  |  |
| DMI (kg/d) | 14.82 | 15.02 | 14.93 | 0.321 | NS |
| Milk Production (kg/d) |  |  |  |  |  |
| Early lactation (0-100 d) | 24.67 | 23.05 | 27.16 | 0.226 | NS |
| Middle lactation (100-200 d) | 17.85 | 17.61 | 15.33 | 0.153 | NS |
| Milk Fat (%) |  |  |  |  |  |
| Early lactation (0-100 d) | 2.96<sup>b</sup> | 3.27<sup>a</sup> | 2.84<sup>b</sup> | 0.093 | ** |
| Middle lactation (100-200 d) | 3.13 | 3.05 | 3.07 | 0.064 | NS |
| Milk Protein (%) |  |  |  |  |  |
| Early lactation (0-100 d) | 2.51<sup>b</sup> | 2.85<sup>a</sup> | 2.71<sup>a</sup> | 0.054 | ** |
| Middle lactation (100-200 d) | 2.93<sup>b</sup> | 3.10<sup>a</sup> | 3.09<sup>a</sup> | 0.068 | ** |
| Total Solids (%) |  |  |  |  |  |
| Early lactation (0-100 d) | 10.95<sup>b</sup> | 11.63<sup>a</sup> | 11.02<sup>b</sup> | 0.178 | ** |
| Middle lactation (100-200 d) | 11.65 | 11.50 | 11.72 | 0.119 | NS |

<sup>†</sup> CG was the control group without supplementation of soybean molasses; WB was the treatments that replaced 15% of corn meal by wheat bran-soybean molasses adsorbent; SH was the treatments that replaced 10% of what bran and 5% corn meal by wheat bran -soybean molasses adsorbent.
<sup>‡</sup>SEM was standard error of means.
<sup>§</sup> NS means the significance among three experimental treatments was not significant ( $P > 0.05$ ).
\* $P < 0.05$ ; \*\* $P < 0.01$ ; \*\*\* $P < 0.001$

#### 2.2.2 Blood biochemistry indexes

The plasma GPT concentration of control was less (*P* < 0.01) than that of WB and SH treatments in early- and mid-lactation periods, while there was no differences (*P* > 0.05) between WB and SH treatments (Table 6). Plasma TP concentration of control was greater (*P* < 0.01) than that of WB and SH treatments in mid-lactation period. The AMY concentration of WB treatment was 96.64% and 32.50% greater (*P* < 0.01) than that of control in early- and mid-lactation periods, respectively, while there was no difference (*P* > 0.05) in AMY concentration between WB and SH treatments. The plasma LDH concentration of WB treatment was 20.87% greater (*P* < 0.01) than that of CG treatments in the mid-lactation period. No differences (*P* > 0.05) in plasma AMM, CHO, GLU, TG and UN concentration were found among three treatments in both early- and mid-lactation periods.

**Table 6.** Effects of different soybean molasses-adsorbents on plasma metabolites in different lactating period in dairy cows

| Item <sup>†</sup> | Group <sup>‡</sup> |  |  | SEM <sup>§</sup> | P <sup>¶</sup> |
| --- | --- | --- | --- | --- | --- |
|  | CG | WB | SH |  |  |
| GPT (U/L) |  |  |  |  |  |
| Early Lactation (0-100 d) | 14.43<sup>b</sup> | 19.00<sup>a</sup> | 19.16<sup>a</sup> | 0.88 | ** |
| Middle Lactation (100-200 d) | 14.64<sup>b</sup> | 19.36<sup>a</sup> | 19.00<sup>a</sup> | 1.65 | ** |
| AMY (U/L) |  |  |  |  |  |
| Early Lactation (0-100 d) | 72.85<sup>b</sup> | 143.25<sup>a</sup> | 112.00<sup>ab</sup> | 23.46 | ** |
| Middle Lactation (100-200 d) | 94.64<sup>b</sup> | 125.40<sup>a</sup> | 109.15<sup>ab</sup> | 19.53 | ** |
| LDH (U/L) |  |  |  |  |  |
| Early Lactation (0-100 d) | 741.67 | 771.56 | 774.50 | 31.00 | NS |
| Middle Lactation (100-200 d) | 702.00<sup>b</sup> | 848.54<sup>a</sup> | 753.93<sup>ab</sup> | 42.61 | ** |
| TP (g/L) |  |  |  |  |  |
| Early Lactation (0-100 d) | 74.78 | 76.38 | 74.88 | 10.25 | NS |
| Middle Lactation (100-200 d) | 81.08<sup>a</sup> | 75.42<sup>b</sup> | 74.26<sup>b</sup> | 12.11 | ** |

<sup>†</sup> GPT = glutamic-pyruvic transaminase; AMY = amylase; LDH = lactate dehydrogenase; TP = total protein;
<sup>‡</sup> CG was the control group without supplementation of soybean molasses; WB was the treatments that replaced 15% of corn meal by wheat bran-soybean molasses adsorbent; SH was the treatments that replaced 10% of what bran and 5% corn meal by wheat bran-soybean molasses adsorbent.
<sup>§</sup> SEM was standard error of means.
<sup>¶</sup> NS means the significance among three experimental treatments was not significant ( $P > 0.05$ ).
\* $P < 0.05$ ; \*\* $P < 0.01$ ; \*\*\* $P < 0.001$

## 3 Discussion

### 3.1 In vitro gas production characteristics of different molasses-adsorbents

*In vitro* maximum gas production is an important parameter to evaluate rumen fermentation in ruminants because it provides valuable information about the kinetics of feed digestion in the rumen and reflects the utilization efficiency of fermentation substrates (Metzler-Zebeli et al., 2012). Khazaal et al. (1993) reported that maximum gas production was positively related to hemicellulose and crude protein (CP) contents, while other studies observed a negative relationship between gas production and CP content of fermentation substrates *in vitro* (Cone and van Gelder, 1999; Tolera and Sundstol, 1999). The current results showed that *V_f_* of SH was greater than that of other the four soybean molasses-adsorbents, due to their differing chemical composition, especially the ratio of non-structural carbohydrate to CP which plays an important role in *in vitro* gas production (Tang et al., 2006).

Indexes of *FRD_0_* and *t_0.5_* usually reflect the rate of degradation at an early incubation stage of < 12 h and the incubation time of reaching half of the maximum gas production, respectively. Generally speaking, the faster *FRD_0_* is, the shorter *t_0.5_* becomes (Wang et al., 2013). In the present study, *FRD_0_* of SH was the least while *t_0.5_* of SH was greatest. These variations of *FRD_0_* and *t_0.5_* should be ascribed to differences of nutrients content among the five soybean molasses-adsorbents.

### 3.2 IVDMD, and IVNDFD of different molasses-adsorbents

*In vitro* DM disappearance can reflect the extent of fermentation of substrates by ruminal microorganisms. Our results showed that IVDMD and IVNDFD of WB were the greatest among the five soybean molasses-adsorbents. It has been shown that dietary molasses supplementation can improve nutrient digestibility in lactating cows, particularly for fiber (Broderick and Radloff, 2004). Usually, dietary sugars undergo rapid fermentation in the rumen of dairy cows, theoretically leading to lactic acid production and decline of ruminal pH, which potentially depresses fiber digestibility (Oelker et al., 2009). However, Broderick and Radloff (2004) reported that replacing high-moisture corn with molasses improved fiber digestibility, likely reflecting a stimulatory effect of molasses on fiber-digesting ruminal bacteria. In the present study, although the adsorbed concentration of soybean molasses was the same, the chemical composition of the molasses-adsorbents differed due to the substrate itself and possibly due to interaction between soybean molasses and the substrate. The rumen is a very complex ecosystem in which numerous microorganisms and factors play an important role in nutrient degradation. Further study is thereby needed to investigate the mechanism of soybean molasses supplementation on the activity of ruminal amylolytic, proteolytic and cellulolytic bacteria during the processes of *in vitro* fermentation.

### 3.3 In vitro fermentation parameters of different molasses-adsorbents

As pH value is an important index reflecting the internal homeostasis of the rumen environment, maintaining a relatively stable ruminal pH is vital to assuring efficient rumen fermentation. Ruminants usually possess highly developed systems to maintain ruminal pH value within a physiological range of about 5.5-7.0 (Krause and Oetzel 2006). In the present study, the pH of *in vitro* incubation fluids ranged from 5.89 to 6.75 for the five soybean molasses-adsorbents. Thus, the highly buffered system maintained suitable conditions for fermentation, microbial growth, and fiber degradation in the rumen (Stewart et al., 1997). Sari et al. (2015) reported that low ruminal pH decreased NH_3_-N concentration and increased non-ammonia N flow compared with high ruminal pH in beef cattle fed diets containing barley straw or non-forage fiber sources. Khalili (1993) found that molasses supplementation linearly decreased the mean value of rumen pH from 6.6 to 6.2 with the increasing levels of molasses fed to crossbred non-lactating cows. However, in our study there was no consistency between NH_3_-N concentration and pH in *in vitro* fermentation fluids, likely because the batch culture system was highly buffered. The inconsistency between in vivo and in vitro results may relate to the buffering capacity of the two systems.

Simultaneously, ruminal NH_3_-N concentration reflects the equilibrium state for CP degradation and synthesis under specific dietary conditions. As an important nitrogen source for microbial growth and protein synthesis, ruminal NH_3_-N has a low efficiency for milk protein synthesis partially due to NH_3_-N losses in the rumen (Tamminga, 1992; Hristov and Ropp, 2003). Satter and Slyter (1974) suggested that the NH_3_-N concentration of rumen fluid should not be less than 5 mg/dL to maintain a suggested that the NH_3_-N concentration of rumen fluid should not be less than 5 mg/dL to maintain a high growth rate of bacteria. Deficiency of NH_3_-N restricts microbial protein synthesis, while high NH_3_-N concentration inhibits the microbial NH_3_-N utilization in the rumen (Hristov et al., 2002). In our study, the NH_3_-N concentrations in *in vitro* fermentation fluids for all five molasses-adsorbents exceeded 5 mg/dL, indicating that the molasses-adsorbents did not restrict ruminal bacterial growth and microbial protein synthesis during the fermentation processes. Meanwhile, feeding a sugar-based product within a diet can change ruminal fermentation pattern, and then further change ruminal NH_3_-N concentration (Broderick and Radloff, 2004; DeFrain et al., 2006). In the present study, the different CP contents of the five molasses-adsorbents probably caused the differences of NH_3_-N concentration in *in vitro* fermentation fluids. In a number of studies, researchers have also observed strong correlation between dietary CP content and NH_3_-N concentration (Broderick and Clayton, 1997). Additionally, the difference in amount and activity of the protein-decomposing microbes for the five substrates might have led to the difference in NH_3_-N concentration. Many studies have demonstrated that protein-decomposing microbes (e.g., *Prevotella sp*.) play an important role in the degradation of CP to NH_3_-N (Jouany, 1996; Wallace, 1996).

### 3.4 In vitro VFA content of different molasses-adsorbents

Ruminal volatile fatty acids (VFAs) are major energy sources for ruminants and differences in the total and proportions of individual VFA are important physiological indices that reflect rumen digestion and metabolism. Ruminal microorganisms can transform carbohydrates (e.g. fiber, starch and soluble sugar) to pyruvic acid, which can be further transferred into different VFAs by metabolic pathways. Several studies have confirmed that molasses addition can reduce the ruminal acetate concentration but increase the ruminal butyrate and propionate concentration *in vitro* and *in vivo* (Hristov and Ropp, 2003; DeFrain et al., 2006; Ferraro et al. 2009). However, Martel et al. (2011) reported that dietary molasses supplementation increased the molar proportions of acetate and butyrate, but decreased the proportions of propionate and total VFA (TVFA) in the rumen of dairy cows. Broderick and Radloff (2004) proposed that increased sugar intake (as dried molasses) does not alter the ruminal concentration of total VFA, acetate, butyrate, or any other individual VFA. In the present study, the concentration of acetate, propionate, butyrate and TVFA of *in vitro* incubation fluids were significantly different for the five soybean molasses-adsorbents, and the largest value of TVFA was obtained for SH, implying that SH may provide more energy for ruminants. Moreover, the variations in VFA concentration might be associated with the differences in ruminal OM digestibility of five molasses-adsorbents (Calsamiglia et al., 2008). Comprehensively considering the *in vitro* fermentation characteristics of the five molasses-adsorbents, especially *in vitro* disappearance of NDF and VFA concentration, two molasses-adsorbents (i.e., wheat bran-molasses, WB; soybean hull-molasses, SH) were selected for further in vivo experiment.

### 3.5 Milk performance

Dietary sugar supplementation can be beneficial for stimulating ruminal microbial protein formation from rumen degradable protein; thus, yield of milk and particularly milk protein content can be easily affected by sugar feeding (Broderick and Radloff, 2004). In the present study, differences in milk production were not observed in early- and mid-lactation, while the milk fat content in early-lactation was improved when dietary corn (100 g/kg) and wheat bran (50 g/kg) were replaced with SH at 150 g/kg of dietary DM, or dietary wheat bran was replaced with 150 g/kg of WB. This result is similar to the previous findings of Martal et al. (2011), who reported that dietary molasses supplementation increased milk fat concentration without significantly affecting milk yield when molasses replaced corn at 50 g/kg dietary DM. However, Brito et al. (2014) reported that yields of milk and milk components can be decreased in lactating cows fed flaxseed meal-based diets supplemented with molasses (liquid molasses plus flaxseed meal vs corn meal plus flaxseed meal). This difference might result from the different molasses sources and dietary composition.

Yan et al. (1997) demonstrated that when molasses inclusion in the diets fed to mid-lactation cows increased from 156 to 468 g/kg DM, milk protein concentration increased from 31.6 to 33.6 g/kg. Keady and Murphy (1998) observed that supplementing sucrose (10 g/kg DM) significantly increased milk protein concentration of lactating dairy cows. The above-mentioned findings support our results that milk protein content was both significantly increased when WB replaced corn at 150 g/kg DM and SH replaced corn at 50 g/kg DM and wheat bran at 100 g/kg DM in early- and mid-lactation periods. Furthermore, Murphy (1999) concluded that milk protein yield can increase when dairy cows are fed rumen-fermentable energy in the form of molasses in a grass silage-based diet. It was suggested that ruminal microbial protein synthesis can be stimulated and a greater proportion of degradable N can be captured by rumen microbes for dairy cows, leading to increased milk protein synthesis. Therefore, the increment of milk protein content in early- and mid-lactation periods is consistent with the previous literature (Broderick et al., 2004).

### 3.6 Blood metabolites

The greater plasma GPT concentration observed when SH replaced corn at 150 g/kg DM and WB replaced corn at 50 g/kg DM and wheat bran at 100 g/kg DM in the dietd fed to cows in early- and mid-lactation might be due to more production of alcohol in the rumen for SH and WB treatments. Generally, *Saccharomyces cerevisiae* populations in the rumen increase when molasses is added to dairy rations, and more alcohol may have been produced by *Saccharomyces cerevisiae* during fermentation of SH and WB treatments. Once in the liver, alcohol casues a rise in plasma GPT concentration (Li et al., 2012; Han et al., 2017).

Dairy cows with high genetic merit require an energy-dense diet to fulfill their production potential, and thus starchy cereals are prevalent in the diets of high-producing dairy cows (Nozière et al., 2014). In the present study, the greater plasma AMY concentration of WB treatment in early- and mid-lactation compared with that of the control treatment probably resulted from molasses being fermented rapidly in the rumen supplying energy to the ruminal microbes leading to a more efficient utilization of starch.

Some studies have shown that LDH can be affected many factors in dairy cows (Chagunda et al., 2006; Piccinini et al., 2007; Wenz et al., 2010). Nyman et al. (2014) confirmed the hypothesis that LDH can indicate inflammation. In the present study, plasma LDH concentration for WB treatment was greater than that of control treatment in mid-lactation. These probably because of the higher formation of lactic acid and lower rumen pH, it resulted in rumen acidosis and caused inflammation.

Lesmeister and Heinrichs (2005) reported no significant differences in blood TP concentration between calves receiving starters containing molasses at 50 and 120 g/kg DM. Azizi-Shotorkhoft et al. (2013) also found no significant difference in blood TP concentration for Moghani sheep fed different levels of molasses (0-100 g/kg). In the current study, the significant changes in plasma TP concentration observed among treatments in mid-lactation, the reasons probably ascrible to the difference in ingredients of experiment diets among treatments. No significant change were observed in early-lactation likely due to the relatively small DMI and a short feeding of the dairy cows during the early lactation period, and the significant differences among treatments during mid-lactation period would be probably resulted from the accumulative effect for a longer feeding.

Thomas et al. (1988) proposed that plasma UN concentration reflects dietary protein intake. Rusche et al. (1993) reported that feeding sources of CP that are less degradable in the rumen decreases blood UN concentration. The lack of significant difference in plasma UN concentration in the present study was in agreement with the findings of Hatfield et al. (1998), who found that molasses type had no effect on plasma UN in sheep.

## 4 Conclusion

Two molasses-adsorbents (soybean molasses adsorbed by wheat bran and soybean hulls) improved maximum gas production, ruminal total VFA content and NDF degradation *in vitro*. Replacement of dietary corn meal/wheat bran by soybean molasses-adsorbents in the diet of dairy cows increased milk protein and fat contents. We conclude dietary corn meal/wheat bran replacement by soybean molasses-adsorbents promotes ruminal fermentation and improves milk quality in lactating dairy cows.

## Acknowledgements

The authors thank Professor Beauchemin Karen for revising the manuscript. This work were supported by the National Natural Science Foundation of China (No.31372342, 3177131431), Youth Innovation Team Project of ISA, Chinese Academy of Sciences (2017QNCXTD_ZCS) and Hunan Provincial Science and Technology Department (2017JJ1028, 2017NH1020).

